# Acetylcholine Enhances Deviance Detection in Hodgkin-Huxley Neuronal Networks

**DOI:** 10.1101/2025.08.11.669609

**Authors:** Fei Fang, Zi-Gang Huang, Zenas C. Chao

## Abstract

The brain’s ability to detect unexpected events, deviance detection (DD), is critical for survival. While DD has been computationally explained by synaptic plasticity, the role of neuromodulators like acetylcholine (ACh) remains less understood. Here, we examine how ACh modulates DD without invoking additional plasticity mechanisms. Using a cholinergic-sensitive Hodgkin-Huxley network of 200 neurons arranged in 2D space and stimulated via five spatially distinct inputs (A–E), we implemented an oddball paradigm with three conditions: standard (80% A, 20% B), deviant (20% A, 80% B), and a multi-standard control (20% each of A–E). ACh levels were controlled by the conductance of the slow K^+^ current. In the absence of ACh, the network already exhibited DD, responding more strongly to deviant A compared to control A. Notably, introducing a small amount of ACh amplified DD, while further increases suppressed it. Maximal DD occurred when strong spike frequency adaptation to standard B reduced competition, and enhanced phase-locking synchronized the network’s response to deviant A. These findings reveal how neuromodulation can shape context-sensitive neural computation, optimizing detection of salient events through a dynamic balance of suppression and synchronization.

## Introduction

Context shapes how sensory information is processed in the mammalian brain. A well-known example is the enhanced neural response to contextually unexpected stimuli, known as deviance detection (DD). This phenomenon, observed even in early sensory cortices, reflects the brain’s ability to detect statistically improbable events or violations of regularities in the environment (Ishishita et al., 2019; Obara et al., 2023; Lao-Rodríguez et al., 2023; Parras et al., 2017; Tiitinen, May, Reinikainen, & Näätänen, 1994; Valerio, Rechenmann, Joshi, De Franceschi, & Barkat, 2024; X.-J. Yu, Xu, He, & He, 2009; McCollum, Manning, Bender, Mendelson, & Anderson, 2024). Critically, DD involves two distinct components: stimulus-specific adaptation (SSA) and genuine deviance detection (genuine DD). SSA refers to reduced neural responses to frequently repeated stimuli, a form of repetition suppression that is specific to stimulus features (Hinz et al., 2025; Valerio et al., 2024; Yarden, Mizrahi, & Nelken, 2022). While SSA can produce larger responses to rare stimuli, this alone does not confirm genuine DD. In contrast, genuine DD requires a stricter criterion: the neural response to a deviant stimulus must exceed the response to the same physical stimulus when it is not surprising (i.e., presented in a control condition). This distinction is essential for isolating the brain’s computation of contextual probability, rather than mere physical novelty.

The capacity for genuine DD is biologically and clinically significant. Impairments in DD have been reported as biomarkers in several neurodevelopmental and psychiatric disorders, including schizophrenia, autism, and attention deficit hyperactivity disorder (De Groote, Bockstael, Botteldooren, Santens, & De Letter, 2021; Näätänen, S. Sussman, Salisbury, & L. Shafer, 2014; Pérez-González, Schreiner, Llano, & Malmierca, 2022), highlighting the need to understand its underlying mechanisms. Genuine DD has been documented across sensory modalities, including auditory, visual, and somatosensory systems (Azouz & Gray, 2003; Bounds & Adesnik, 2025; Derveer, Ross, & Hamm, 2023; McCollum et al., 2024; Obara et al., 2023; Parras et al., 2017; Wang, Rangarajan, Gerfen, & Krauzlis, 2018), and across species ranging from rodents to humans (Behrens, Bäuerle, Kössl, & Gaese, 2009; Ulanovsky, Las, Farkas, & Nelken, 2004; Yarden & Nelken, 2017). It has also been observed throughout the brain, from subcortical structures like the inferior colliculus to higher cortical areas (Chen, Helmchen, & Lütcke, 2015; Tsolaki, Kosmidou, Hadjileontiadis, Kompatsiaris, & Tsolaki, 2015). Notably, recent work has demonstrated genuine DD in neuronal cell cultures (Zhang et al., 2025), showing that this computation can emerge from simple networks, independent of complex brain circuitry.

Computational models have typically attributed genuine DD to synaptic plasticity. While models based on long-term plasticity exist (Hertäg & Sprekeler, 2020; Wacongne, Changeux, & Dehaene, 2012), the rapid emergence of DD *in vivo* points to fast-acting mechanisms. Short-term synaptic plasticity, particularly short-term depression, has been a primary candidate (Anwar, Li, Bucher, & Nadim, 2017; Mill, Coath, Wennekers, & Denham, 2011; Yarden & Nelken, 2017). More recent studies show that intrinsic neuronal adaptation, such as threshold adaptation, is also sufficient to generate genuine DD. Moreover, short-term depression and threshold adaptation can interact synergistically, enhancing DD beyond what either mechanism achieves alone (Kern & Chao, 2023). These findings suggest that genuine DD can arise from the dynamic interaction of fast plasticity mechanisms, without requiring long-term structural changes.

Beyond synaptic and intrinsic mechanisms, neuromodulation also plays a key role in shaping neural responsiveness (Brzosko, Mierau, & Paulsen, 2019; Ghosh & Maunsell, 2024; Sarter & Parikh, 2005; Shine et al., 2021; Thiele & Bellgrove, 2018; Wiseman et al., 2024; A. J. Yu & Dayan, 2005). In particular ACh modulates neuronal excitability and synaptic transmission, and has been implicated in DD (Hasselmo & McGaughy, 2004; Herrero et al., 2008; Kuczewski, Aztiria, Gautam, Wess, & Domenici, 2005; Pérez-González, Lao-Rodríguez, Aedo-Sánchez, & Malmierca, 2024; Pérez-González et al., 2022; Picciotto, Higley, & Mineur, 2012). ACh enhances neural gain and modulates inhibition by acting on presynaptic interneurons (Colangelo, Shichkova, Keller, Markram, & Ramaswamy, 2019). It also influences short-term synaptic plasticity via muscarinic and nicotinic receptors that regulate presynaptic transmitter release (McKay, Placzek, & Dani, 2007). Functionally, ACh increases the precision of bottom-up sensory transmission (Kang, Huppé-Gourgues, & Vaucher, 2014), which is essential for predictive coding, particularly in tuning the gain on prediction error signals (Pérez-González et al., 2024). Notably, ACh has been shown to modulate genuine DD without affecting SSA (Moran et al., 2013). However, the exact neural mechanisms by which ACh contributes to genuine DD remain unclear.

To address this, we constructed a cholinergic-regulated Hodgkin-Huxley (HH) network model devoid of other forms of synaptic plasticity. We systematically varied ACh levels across the network during a simulated oddball paradigm. Our results show that genuine DD is maximized at an intermediate level of ACh. Mechanistically, this effect arises from two complementary processes: local suppression via ACh-induced spike frequency adaptation (SFA), which frees up neural resources for deviant detection, and global synchronization via ACh-induced phase locking, which amplifies the network’s response to the deviant stimulus.

## Results

### A 2D network of cholinergic Hodgkin-Huxley spiking neurons

We used a HH-based model of cholinergic modulation in pyramidal cells to simulate neuron membrane potential dynamics (Hodgkin & Huxley, 1952; Roach, Eniwaye, Booth, Sander, & Zochowski, 2019). Cholinergic signaling through M1 muscarinic ACh receptors can be well modeled by parameterizing the maximal conductance *g*_*ks*_ of a slow, low-threshold potassium-mediated adaptation current (see details in Methods).

To characterize the fundamental properties of synaptic transmission within our model, we first simulated a minimal two-neuron excitatory circuit (Figure 1A). The presynaptic neuron (Neuron 1) was stimulated with a brief, suprathreshold current pulse (*I*_*drive*_) at the beginning of the simulation. This single stimulus was sufficient to elicit a stable, periodic train of action potentials in Neuron 1 for the entire 500 ms recording period (Figure 1B, *V*_1_). The activity of Neuron 1 was effectively and reliably propagated to the postsynaptic neuron (Neuron 2). We observed a precise one-to-one firing relationship, where each action potential in the presynaptic neuron triggered a corresponding action potential in the postsynaptic neuron with a minimal delay (Figure 2B, *V*_2_). This faithful transmission was mediated by the dynamics of the excitatory synapse. Following each presynaptic spike, the postsynaptic excitatory conductance (*g*_*e*_) was transiently activated, leading to a sharp, inward synaptic current (*I*_*syn*_) in Neuron 2. These excitatory postsynaptic currents consistently depolarized the postsynaptic membrane to its firing threshold, thus ensuring robust signal transmission.

**Figure 1.**
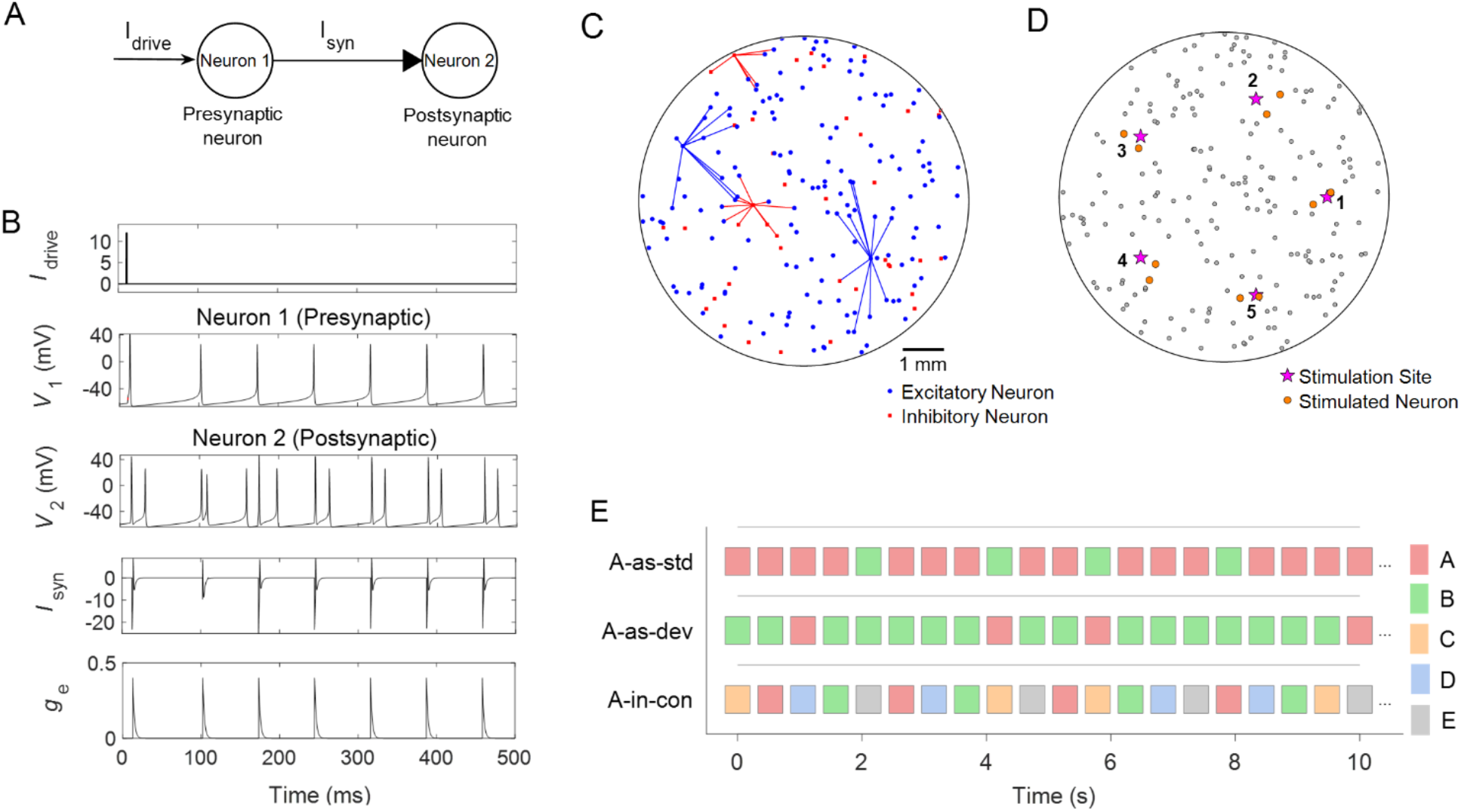
Neural network model and experimental setup. (**A**) Schematic of a two-neuron excitatory circuit. *I*_*drive*_ and *I*_*syn*_ represent external and synaptic currents, respectively. (**B**) Responses to a brief pulse of current stimulation. *V*_*1*_ and *V*_*2*_ denote the membrane potentials of Neurons 1 and 2 in panel A. The excitatory synaptic conductance is denoted as *g*_*e*_. (**C**) Example network layout, illustrating the spatial locations of excitatory (red) and inhibitory (blue) neurons, along with all outgoing synaptic connections from two neurons of each type. (**D**) Stimulation design. Five stimulation sites (A–E) are evenly spaced on a circle with a 2.5 mm radius. When a site is stimulated, current is applied to the two neurons nearest to the electrode (highlighted in yellow). (**E**) Oddball stimulation paradigm. Each square represents a stimulation event, with the stimulus electrode indicated by color. Three sequence types are shown: A-as-std, A-as-dev, and A-as-con. Note that stimuli were delivered instantaneously (1 ms duration); the extended durations in the figure are for visualization only.

**Figure 2.**
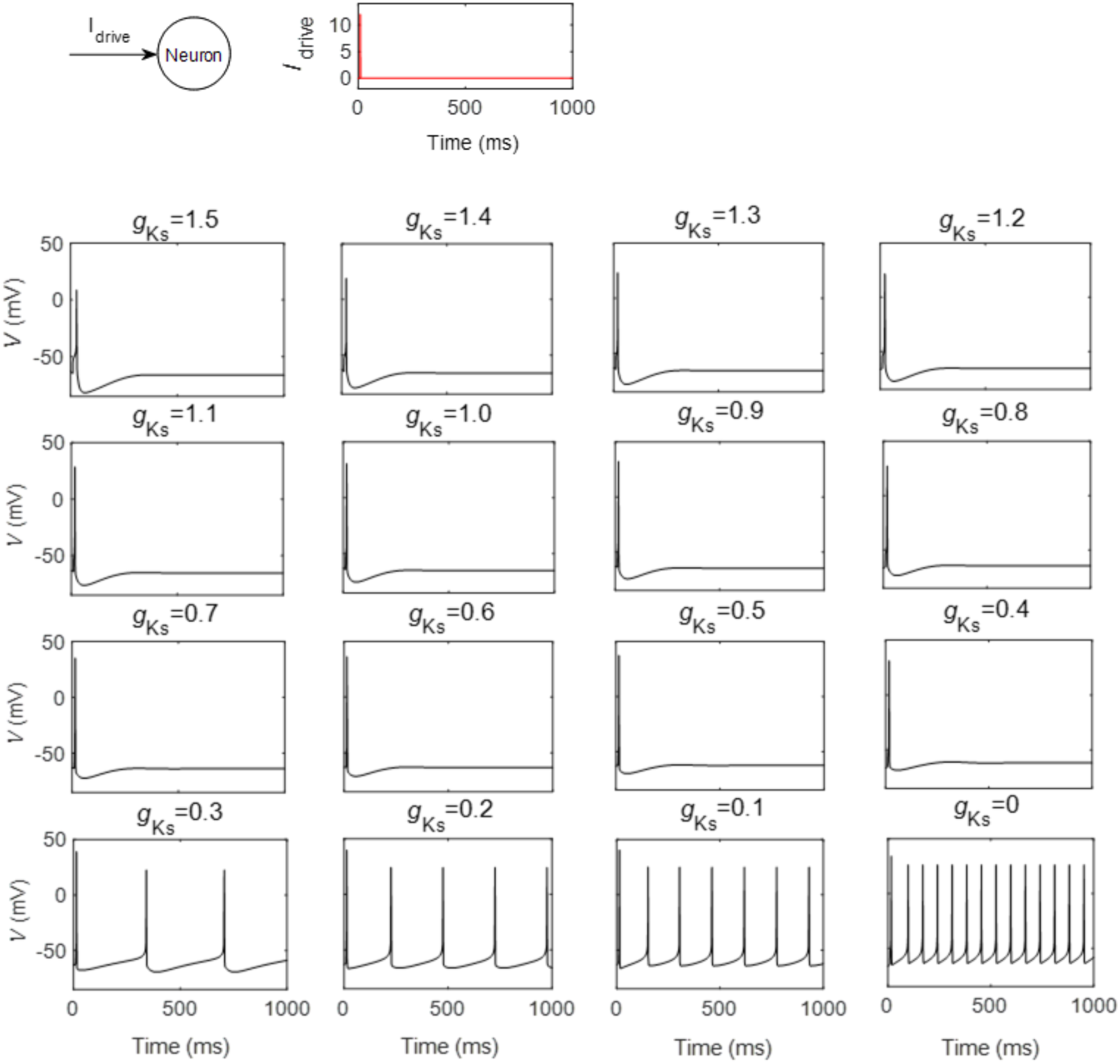
Effect of slow potassium conductance (*g*_*ks*_) on the firing pattern of a single HH neuron. Schematic of the setup: a single HH neuron receives a brief external current input (*I*_*drive*_). Each panel shows the neuron’s membrane potential under a different *g*_*ks*_ value, ranging from 1.5 (corresponding to no ACh) to 0 (representing maximal ACh).

We constructed a network of 200 HH neurons (Figure 1C), consisting of 160 excitatory (80%) and 40 inhibitory (20%) neurons. These neurons are randomly distributed within a two-dimensional circular area with a radius of 4 mm, resembling a neuronal cell culture (Zhang et al., 2025) Each neuron forms synapses with 10 randomly selected postsynaptic neurons within a distance of 2 mm for excitatory neurons and 1 mm for inhibitory neurons. The synaptic weight is set at w = 1.0 nS (see details in Methods). These specific parameters were chosen to ensure reliable responses to strong stimuli without inducing unstable or persistent activity. Connectivity and weights remain constant throughout the simulation. Stimulation is applied through five “virtual electrodes” (indicated by stars in Figure 1D), evenly spaced at a distance of 2.5 mm from the center of the dish. These correspond to stimuli A, B, C, D, and E. Upon stimulation, current is injected into the three neurons (either excitatory or inhibitory) closest to the stimulus site (highlighted in yellow in Figure 1D).

It is important to note that our goal is to isolate the effect of ACh; therefore, we use the simplest form of network dynamics, excluding both short-term plasticity (e.g., neuronal threshold adaptation or synaptic depression) and long-term plasticity (e.g., spike-timing-dependent plasticity).

### Oddball paradigm and simulation

To investigate genuine DD, we used a classical oddball paradigm (Figure 1E), presenting a total of 200 stimuli at regular intervals of 100ms (see details in Methods). The target stimulus (labeled A) and distractor stimuli (labeled B, C, D, and E) were presented in three randomized sequences: (1) the standard sequence (A-as-std), consisting of 160 presentations of A and 40 of B; (2) the deviant sequence (A-as-dev), with 40 presentations of A and 160 of B; and (3) the many-standards control sequence (A-in-con), with 40 presentations of each stimulus (A–E). All stimuli were presented in random order. Each stimulus consisted of a 12 *μA*/*cm*^2^ current pulse lasting 1 ms. In this design, stimulus A serves three roles: (1) as an expected stimulus in the standard sequence (A-as-std), (2) as an infrequent violation of the expected stimulus B in the deviant sequence (A-as-dev), and (3) as an equally infrequent stimulus in the many-standards control sequence (A-in-con), where no strong expectation is formed, allowing control for pure adaptation effects.

We constructed five networks with different structures. For each network, we applied the three stimulation sequences, each repeated four times with a different random stimulation order. A 1-second stabilization period was inserted between sequences to allow the network to fully recover before the next sequence began.

### Cholinergic effect on HH neuron and network

We first examined how the slow potassium current influences the firing properties of a single HH neuron by systematically varying *g*_*ks*_, the maximal conductance of the slow potassium current. As illustrated in Figure 2, the neuron was subjected to a suprathreshold depolarizing current (*I*_*drive*_) to elicit action potentials. When *g*_*ks*_ is high or intermediate (e.g., 1.5 to 0.4 mS/cm^2^), the neuron fires a single action potential, after which the membrane potential repolarizes and remains near the resting level (Figure 2). As *g*_*ks*_ is progressively decreased, reflecting increasing levels of ACh, the firing pattern shifts. For *g*_*ks*_ ≤ 0.3, the neuron transitions to a tonic firing mode, marked by sustained, rhythmic action potentials throughout the stimulus duration. These results demonstrate that the slow potassium current is a key mechanism for regulating the transition between phasic and tonic firing modes in single neurons.

Figure 3A shows the spiking responses of a sample network of 200 HH neurons to the three stimulation sequences under two levels of ACh. At *g*_*ks*_ = 0.3 (high ACh), the network exhibited strong but non-time-locked responses. In contrast, at *g*_*ks*_ = 1.3 (low ACh), evoked responses were less frequent but more tightly time-locked to the stimuli. Examples membrane potentials of individual neurons are shown in Figure 3B.

**Figure 3.**
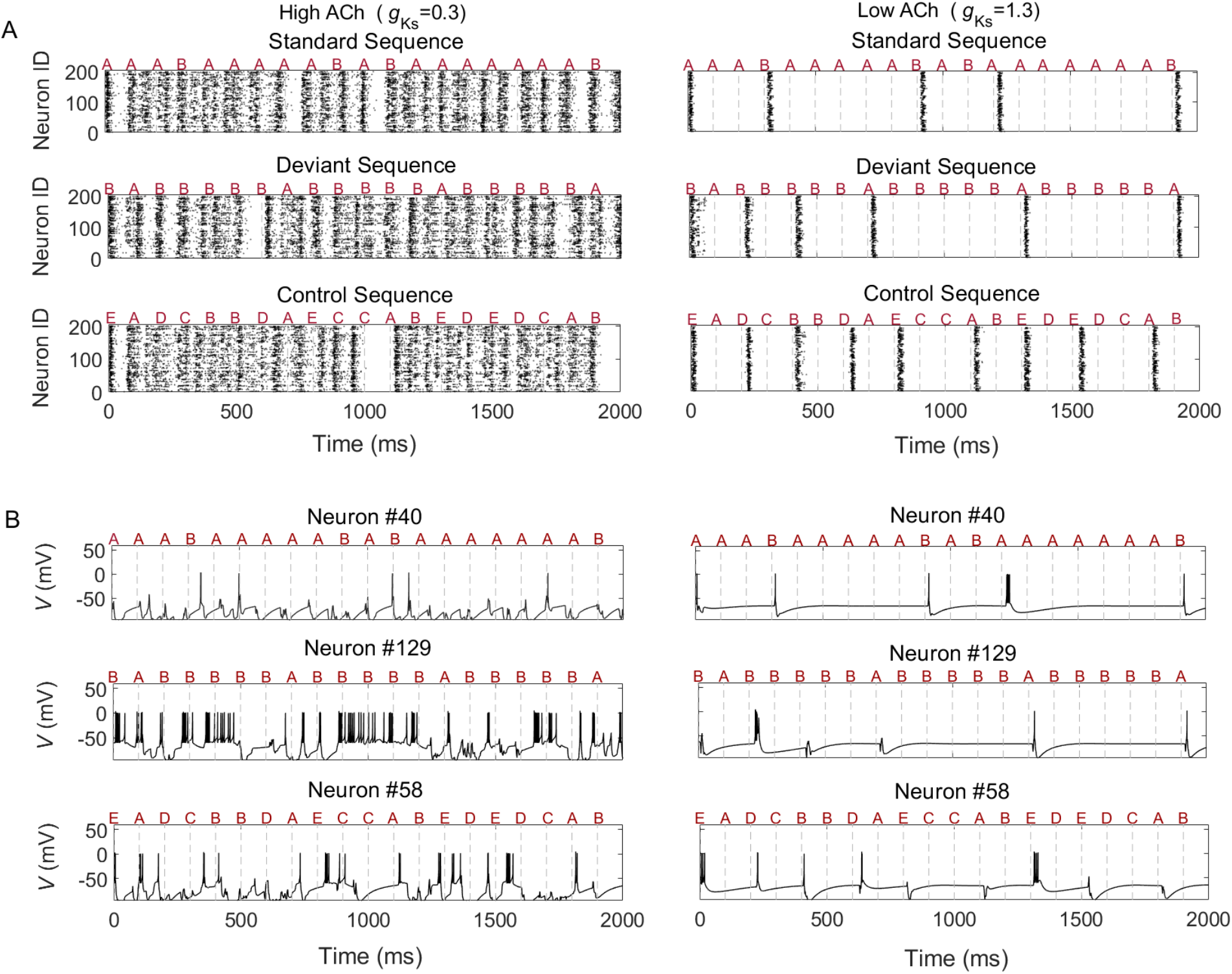
Network activity under different levels of *g*_*ks*_. (**A**) Raster plots showing the spiking activity of all 200 neurons in the network under high ACh (low *g*_*ks*_ = 0.3) and low ACh (high *g*_*ks*_ = 1.3) conditions. Different stimulation types (A–E) are indicated by red letters above the plot. Sequence types vary across rows. (**B**) Membrane potentials of three example neurons (#40, #129, and #58) from the network shown in panel A.

### Cholinergic modulation of genuine deviance detection

To demonstrate that our model exhibits genuine DD, we first examined the evoked responses to stimulus A across all three sequences (A-as-dev, A-in-con, and A-as-std) in a sample network under four different *g*_*ks*_ values (1.5, 1.3, 0.8, and 0.3 mS/cm^2^) (Figure 3). When *g*_*ks*_ = 1.5 (no ACh), stimulus A in the deviant sequence elicited a robust response with a latency of approximately 28.5 ms. In contrast, responses to A in the control sequence were less consistent, and in the standard sequence, only sporadic activity was observed. At *g*_*ks*_ = 1.3 (low ACh), strong evoked responses were still observed, but the latency shifted earlier to around 15.5 ms. When *g*_*ks*_ was reduced further to 0.8, evoked responses began to emerge even in the standard sequence, and the difference in response rates between the deviant and control sequences diminished. Finally, at *g*_*ks*_ = 0.3 (high ACh), evoked responses became weaker across all sequences, and late, scattered responses were more frequently observed. These response patterns were further quantified using post-stimulus time histograms, which measure the average number of spikes per millisecond across trials for each sequence (Figure 3B).

We then calculated the deviance detection index (DDI), which quantifies the relative response magnitude between the deviant and control sequences (see Methods), in the sample network across different *g*_*ks*_ values ranging from 1.5 to 0 (Figure 4C). When ACh was absent (*g*_*ks*_ = 1.5), the DDI was low (0.17) but not zero, suggesting that the HH neural network, despite lacking any form of plasticity, is still capable of detecting deviant stimuli (see Discussion for further details). Introducing a small amount of ACh (*g*_*ks*_ = 1.3) increased the DDI to a peak of 0.361. As *g*_*ks*_ was further reduced, the DDI gradually declined to zero, indicating that the network responses to the deviant and control sequences became increasingly similar (as shown in Figure 4A).

**Figure 4.**
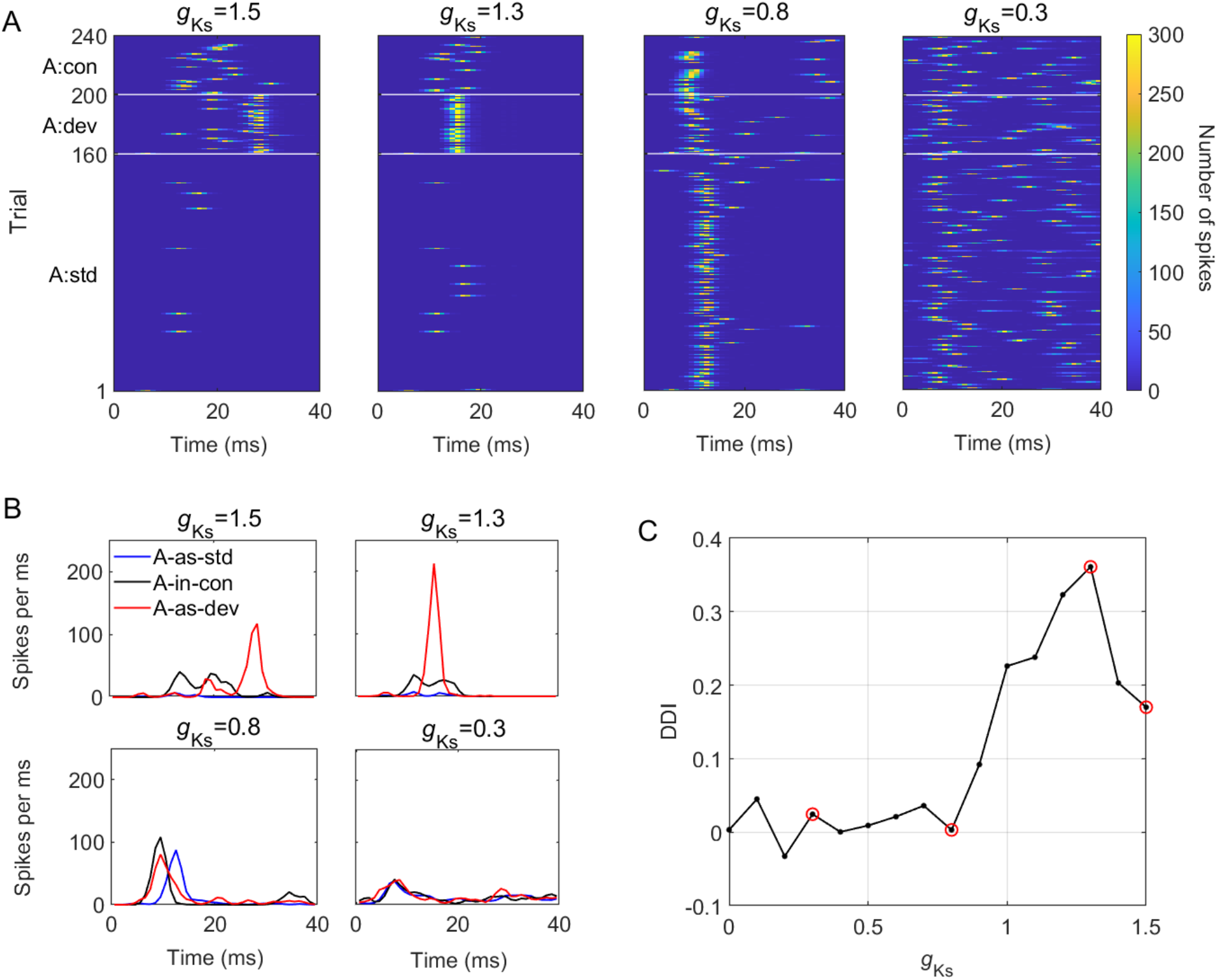
Deviance detection in a sample network under varying *g*_*ks*_ levels. (**A**) Network responses to the target stimulus A under three sequence types. A:std, A:dev, and A:con represent the evoked responses to stimulus A under the A-as-std (160 trials), A-as-dev (40 trials), and A-as-con (40 trials) sequences, respectively. Color indicates the total spike count across all neurons per millisecond. Four *g*_*ks*_ conditions (1.5, 1.3, 0.8, and 0.3) are shown. (**B**) Post-stimulus time histograms for network responses to stimulus A under the three sequence types. (**C**) DDI for each *g*_*ks*_ value in the sample network. The four examples shown in panels A and B are marked with red circles.

To ensure that the results were not dependent on the specific choice of stimulus sites or network structures, we repeated the simulation using four different stimulation sites across five different networks. This yielded a total of 20 simulation sets, each with *g*_*ks*_ values ranging from 0 to 1.5. The network responses for the A-as-dev and A-in-con sequences are shown in Figure 5A, and the corresponding 20 DDI vs. *g*_*ks*_ curves are presented in Figure 5B. To further assess the cholinergic effect on DDI, we compared DDI values across different *g*_*ks*_ levels to those at *g*_*ks*_ = 1.5 (no ACh) (pairwise one-tailed t-tests). The results showed that DDI was significantly higher at *g*_*ks*_ values of 1.4, 1.3, and 1.2 (Figure 5B). These comparisons are further illustrated in Figure 5C. These results confirm that a small amount of ACh maximizes deviance detection.

**Figure 5.**
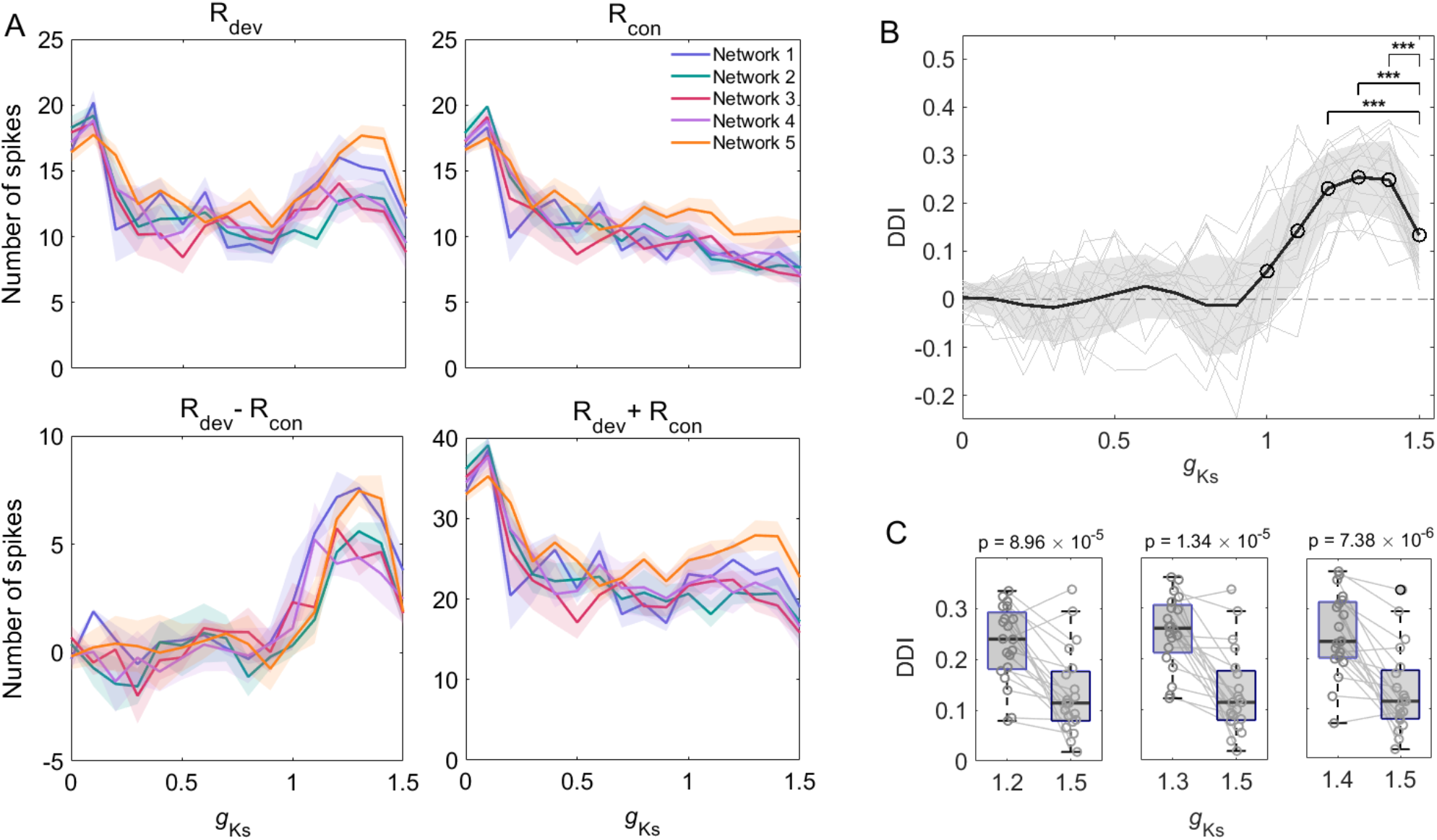
Cholinergic modulation of deviance detection. (**A**) Average network responses to stimulus A in the A-as-dev (R_dev_) and A-as-con (R_con_) sequences. The difference (R_dev_ – R_con_) and sum (R_dev_ + R_con_) are also shown. The DDI is defined as the ratio of the difference to the sum. (**B**) DDI values plotted against *g*_*ks*_ across 20 simulations (5 network configurations × 4 stimulation sites). The black line indicates the average trend. DDIs significantly greater than zero (α = 0.05, one-tailed Wilcoxon test with Benjamini–Hochberg method) are marked with black circles. Asterisks indicate values significantly greater than the no-ACh condition (*g*_*ks*_ = 1.5). (**C**) Conditions where DDI values (*g*_*ks*_ = 1.2, 1.3, and 1.4) were significantly higher than at *g*_*ks*_ = 1.5. Corresponding p-values are shown above each comparison.

### Cholinergic modulation of local suppression via spike frequency adaptation

In this section, we aim to provide a mechanistic explanation of the observed cholinergic facilitation of DD. We first validated an existing theory of genuine DD proposed by Kern et al. (Kern & Chao, 2023), which suggests that either neuronal threshold adaptation or synaptic depression—two forms of short-term plasticity—reduces the propagation of evoked responses to standard stimuli (in this case, stimulation B). This suppression frees up neural resources for responding to deviant stimuli (stimulation A). Such local suppression is less prominent in the control sequence, where stimuli are spatially distributed, which together contributes to a higher DDI. Although our model does not include explicit mechanisms for neuronal threshold adaptation or synaptic depression, SFA can be induced by ACh in HH neurons (Cui & Strowbridge, 2019; Roach et al., 2019).

Figure 6A illustrates the results of SFA. An external current pulse (500 ms duration) was applied to a neuron with *g*_*ks*_ = 0.1 and to one with *g*_*ks*_ = 1.5 (representing no ACh). At *g*_*ks*_ = 0.1, spiking activity was sustained with minimal changes in inter-spike intervals (ISIs). In contrast, at *g*_*ks*_ = 1.5, the firing rate was reduced (as shown previously in Figure 2), and the ISIs progressively lengthened, indicating SFA. This adaptation can be quantified by the degree of ISI change over time (see Methods and (Roach et al., 2019)). As shown in Figure 6B, SFA depends on the magnitude of the input current. In general, stronger input currents produce weaker SFA, whereas higher *g*_*ks*_ values lead to stronger SFA. Note that in some cases, particularly under very strong stimulation, SFA does not occur, as indicated by the white regions in Figure 6B. This occurs because, under strong and sustained current, the neuron fires only a few spikes before the membrane potential becomes trapped at a highly depolarized level (e.g., −20 mV), leading to sodium channel inactivation and preventing further spiking (see examples in Supplemental Figure S1). Therefore, we defined the current range in which SFA occurs as Δ*I*_*drive*_, which estimates how likely synaptic currents in the network are to induce SFA (indicated in Figure 6B). In other words, networks with larger Δ*I*_*drive*_ values are more likely to exhibit strong SFA in postsynaptic neurons due to synaptic transmission. When Δ*I*_*drive*_ is plotted against *g*_*ks*_ (Figure 6C), the maximum value of 23 is observed at *g*_*ks*_ = 1.25, around the same condition under which the highest DDI is observed. This result supports the idea that ACh modulates DDI through SFA, a local suppression mechanism that limits the further propagation of activity.

**Figure 6.**
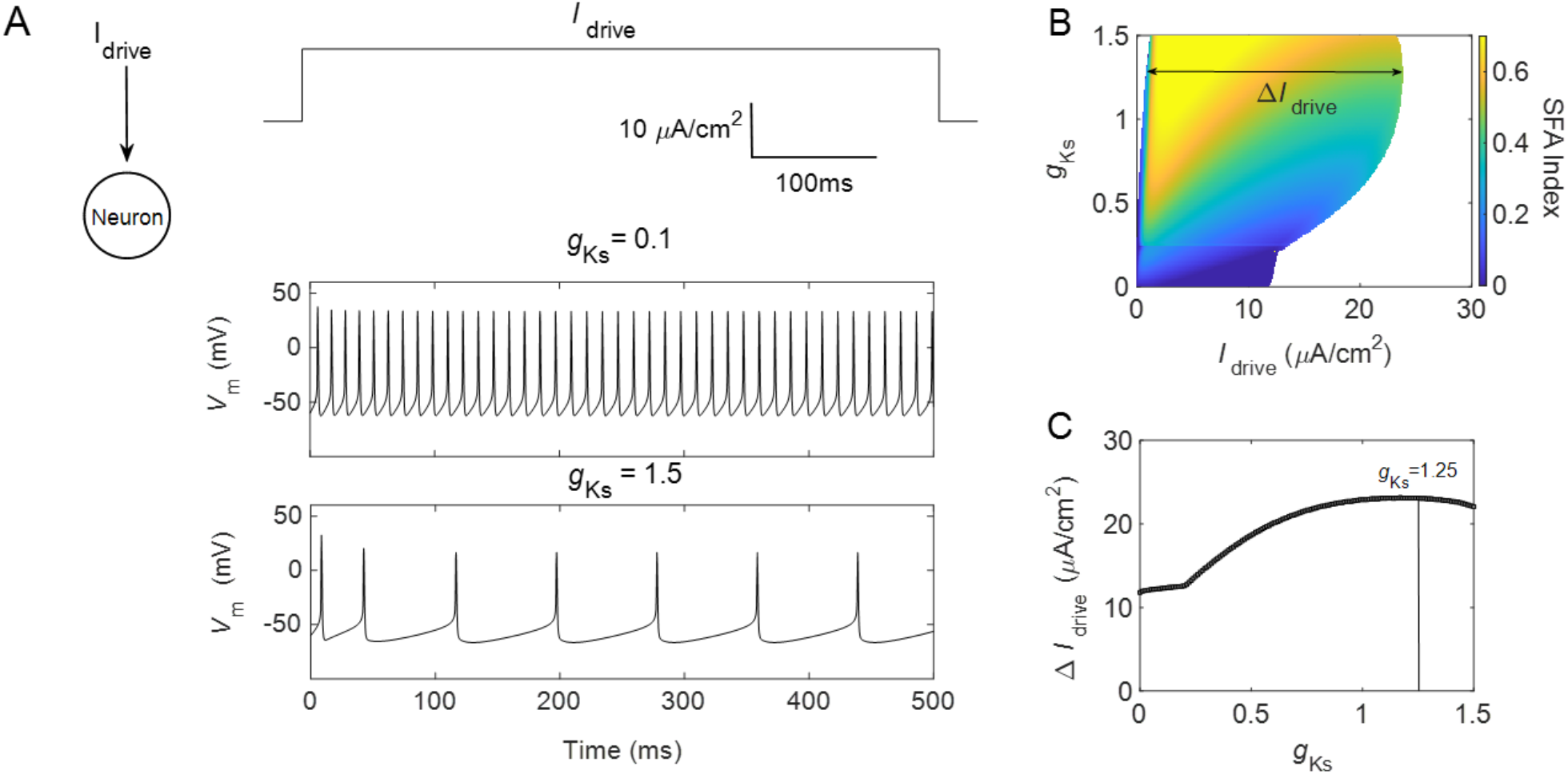
Cholinergic modulation of spike frequency adaptation. (**A**) Responses of a HH neuron to sustained current stimulation under different *g*_*ks*_ values. (**B**) SFA as a function of *g*_*ks*_ and stimulation current (*I*_*drive*_). White areas indicate conditions where SFA is absent (see main text for details). For each *g*_*ks*_ level, Δ*I*_*drive*_ is defined as the range of current over which SFA occurs. (**C**) Δ*I*_*drive*_ as a function of *g*_*ks*_. The maximum Δ*I*_*drive*_ is observed at *g*_*ks*_ = 1.25 (indicated by the vertical line).

### Cholinergic modulation of global synchronization via phase locking

In addition to local suppression, cholinergic modulation can also promote phase locking, where neuronal firing becomes more synchronized (Roach et al., 2019). We quantified phase locking using mean phase coherence (MPC), which reflects the average synchrony between pairs of neurons, ranging from 0 (random firing) to 1 (perfect phase locking). Figure 7 shows MPC values across different *g*_*ks*_ levels for all five networks under both deviant and control sequences. When *g*_*ks*_ > 0.5, MPC for the deviant sequence becomes greater than that for the control sequence, reaching a maximum at *g*_*ks*_ = 1.3, where MPC peaks at 0.64. This result suggests that the network’s ability to synchronize is maximized at *g*_*ks*_ = 1.3 under the deviant sequence. At this point, the probability of neurons firing simultaneously increases, enhancing the response to deviant stimuli. This, in turn, amplifies the difference between responses to deviant and control sequences, leading to a maximal DDI.

**Figure 7.**
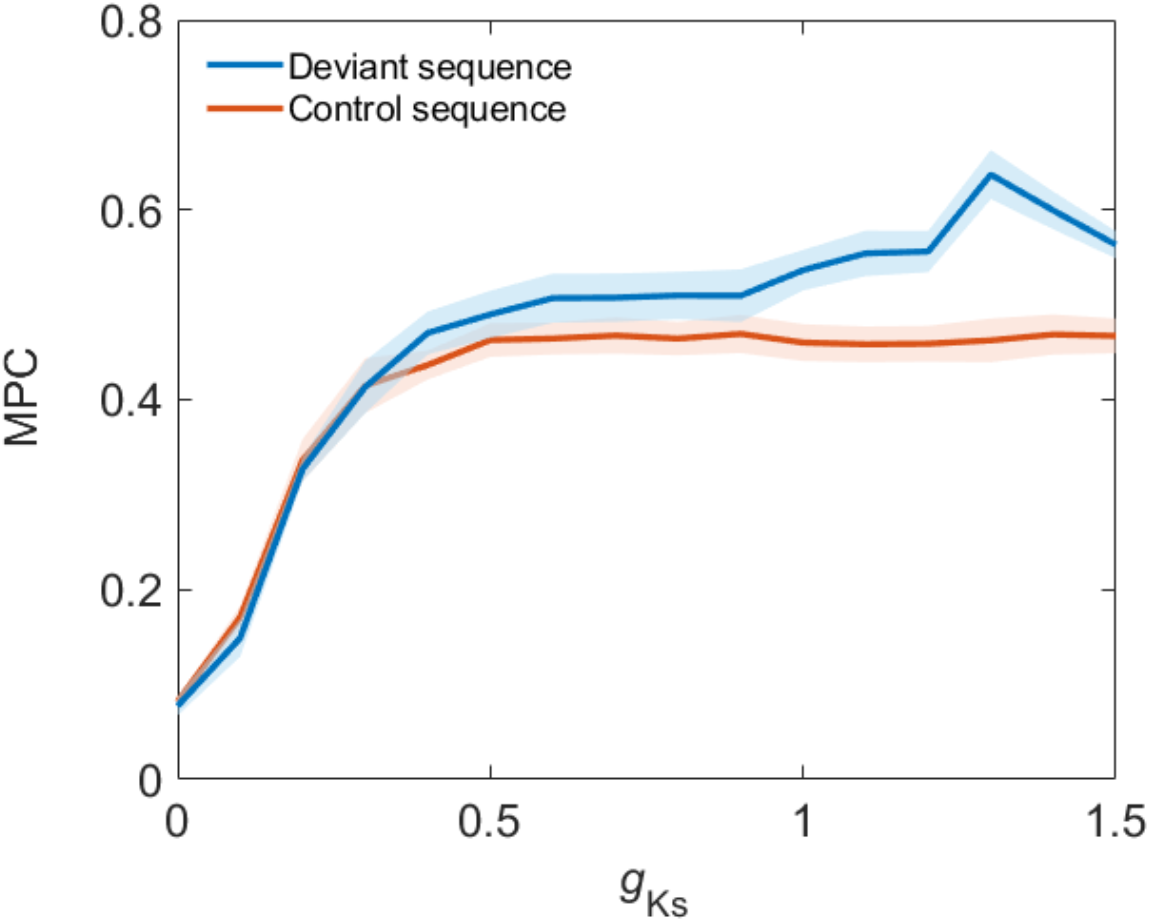
Cholinergic modulation of phase-locking in the network. Mean phase coherence (MPC), a measure of average synchrony between pairs of neurons, was computed for the A-as-dev and A-as-con sequences under different *g*_*ks*_ values. MPC was calculated across all neuron pairs and 5 simulations (5 network configurations). The mean and standard deviation are shown.

Together, we propose that at *g*_*ks*_ = 1.3, the combination of maximal local suppression, driven by the strongest SFA to the standard stimulus B, which frees the greatest amount of neural resources for the deviant stimulus A in the A-as-dev sequence, and the strongest globally synchronized response to A due to maximal phase locking, results in the most robust evoked response to A. This interplay results in the maximal DDI (see illustration in Figure 8).

**Figure 8.**
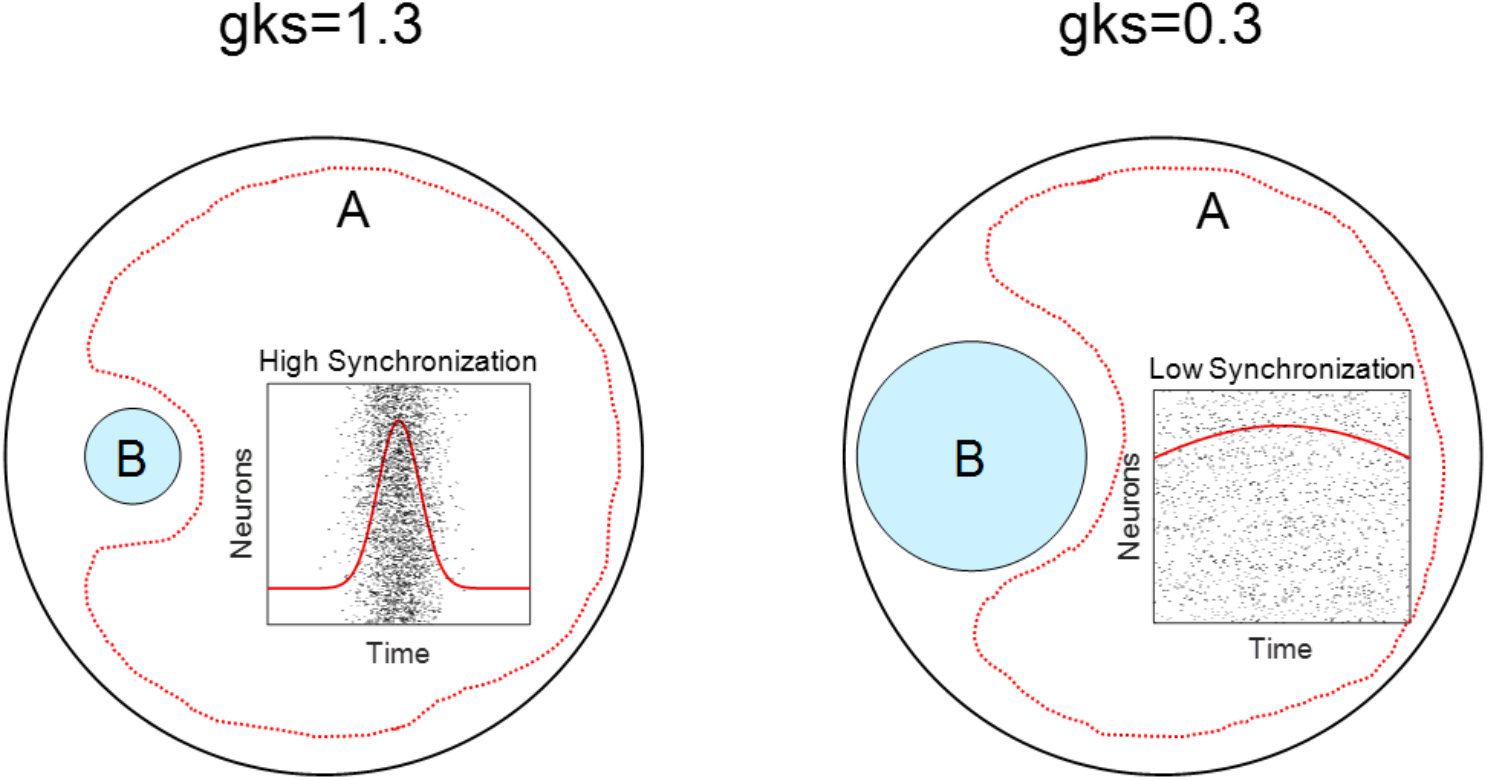
Conceptual model of cholinergic modulation in genuine DD. In the A-as-dev sequence, when *g*_*ks*_ = 1.3 (left), spike frequency adaptation is strongest. As a result, repeated standard stimulation (stimulus B) suppresses activity near the electrode, preventing further propagation of evoked responses (indicated by the small blue circle). This suppression frees up the rest of the network (red dotted area) to respond more robustly to the deviant stimulus A. In addition, phase-locking is maximal at *g*_*ks*_ around 1.3, resulting in stronger synchronization and a more prominent network-wide response to A. In contrast, when *g*_*ks*_ deviates from 1.3 (e.g., 0.3; right panel), both local suppression and global synchronization are reduced, leading to a weaker deviant response.

## Discussion

We use a HH neural network to show that cholinergic modulation can suppress neuronal firing and enhance interneuronal synchronization, enabling genuine DD. This finding indicates that spatially distributed neural computation allows the network to retain a memory of both the stimulus identity and its regularity, and suggests that genuine DD can arise through purely feedforward signaling, without the need for top-down modulation and long-term plasticity.

### Mechanisms of genuine deviance detection

Our results show that even when *g*_*ks*_ is set to 1.5, representing a low ACh state, DDI is 0.17, indicating a residual ability to detect deviations (see Figure 3). This effect can be attributed to SFA, which remains active at *g*_*ks*_ = 1.5 (Cui & Strowbridge, 2019; Roach et al., 2019). SFA is a non-synaptic plasticity mechanism that reduces neuronal firing rates over repeated stimulation. It operates similarly to threshold adaptation (Fontaine, Peña, & Brette, 2014; Shaban, Bezugam, & Suri, 2021), but while threshold adaptation raises the spiking threshold, SFA reduces spike counts by lowering discharge frequency. Kern & Chao (Kern & Chao, 2023) demonstrated that threshold adaptation alone is sufficient for DD. They proposed a dual mechanism of local fatigue and global recovery: neurons near the repeated standard stimulus become suppressed (local fatigue), which prevents widespread activation. As a result, the rest of the network remains responsive (global recovery), allowing strong responses to unexpected stimuli. Our cholinergic HH model replicates this mechanism: ACh-regulated SFA induces local fatigue around repetitive stimuli, preserving broader network sensitivity for detecting deviants.

Kern and Chao (2023) reported a synergistic effect between threshold adaptation (functionally equivalent to SFA in our model) and short-term synaptic depression. Both neuronal and synaptic forms of short-term plasticity can support genuine DD, and their effects can be additive. To explore this, we incorporated short-term synaptic depression into our model (see Methods for model details) and observed a further enhancement of DD (Figures S2 and S3). The difference in DDI with and without short-term depression is shown in Figure S4. For example, at *g*_*ks*_ = 1.3, the DDI increased from 0.316 with ACh alone to 0.410 when both ACh and short-term depression were present. We propose that any short-term adaptive mechanism operating across space can encode information about both stimulus identity and its regularity, thereby supporting genuine DD. This theory does not preclude a role for long-term plasticity in genuine DD (Azouz & Gray, 2003; Bounds & Adesnik, 2025; Derveer et al., 2023; Obara et al., 2023; Parras et al., 2017).

### Synchronization and latency shifts: temporal dynamics of deviance detection

We found that network synchronization was highest at *g*_*ks*_ = 1.3 in the deviant sequence (Figure 7), which corresponds to a state of low ACh and Type 2 neuronal excitability. According to the mechanisms described by Roach et al. (Roach et al., 2019), Type 2 neurons possess two key properties that facilitate synchronization: 1) a flat frequency-current (f/I) curve (see f/I curves of our model under different *g*_*ks*_ values in Figure S5), which ensures firing rates are uniform across the network, and 2) a biphasic phase-response curve (see phase-response curves of our model under different *g*_*ks*_ values in Figure S6), which allows neurons to both advance and delay their spike times to align with each other. At *g*_*ks*_ = 1.3, these properties yield a latency of 15.5 ms, not the latency of single neurons, but the timing of a collective, synchronous burst characteristic of a phase-coding regime. When *g*_*ks*_ decreased to 1.0 (see Figure S7), latency advanced to 11.5 ms, consistent with a shift to rate coding (Type 1 excitability). In this high-ACh state, the neuronal f/I curve becomes steeper, producing high gain: small inputs trigger a large, rapid increase in firing rate. The network then shifts from encoding information in the precise timing of spikes within a synchronized population to encoding it in the firing rate of individual neurons. Thus, *g*_*ks*_ functions as a dynamic control parameter, switching the network between two computational regimes: phase coding with synchronized bursts versus rate coding with rapid, individual responses, and in doing so, modulates the temporal properties of DD.

### Biological relevance of *g*_*Ks*_ and cholinergic modulation

We showed that moderate ACh levels enhance DD (Figures 3 and 4). In our model, cholinergic modulation of neuronal excitability is implemented by varying the maximal conductance of the M-current (a slow, muscarinic receptor-regulated potassium current) represented by the parameter *g*_*ks*_. This approach is grounded in well-established findings that ACh suppresses the M-current, thereby increasing neuronal excitability (Brown & Adams, 1980; McCormick & Prince, 1985). High ACh levels, which strongly block the M-current, are modeled as a low conductance value (*g*_*ks*_ = 0), producing Type 1 excitability. Conversely, low ACh levels, where the M-current remains active, are modeled as a high conductance (*g*_*ks*_ = 1.5), producing Type 2 excitability (Roach et al., 2019). This parameterization is quantitatively supported by the concentration-response relationship reported by Adams et al. (Adams, Brown, & Constanti, 1982), who found that the ACh agonist muscarine inhibits the M-current in a dose-dependent manner, with near-maximal suppression (∼80%) at ≥10 μM and a half-maximal effective concentration (EC_50_) of ∼3 μM. Thus, in our model, the *g*_*ks*_ = 0 condition could correspond directly to the experimentally observed state of near-complete M-current blockade under high cholinergic tone (≥10 μM muscarine). In future work, we will use the estimated concentration to test the cholinergic effects on genuine DD in neuronal culture.

### Spatial heterogeneity of ACh and implications for modeling

Our current model assumes a homogeneous distribution of ACh by using the same *g*_*ks*_ value across all neurons in the network. While earlier models similarly treated ACh modulation as spatially diffuse (Li et al., 2018; Picciotto et al., 2012), more recent studies highlight its spatially localized and dynamic nature (Lohani et al., 2022; Zaborszky et al., 2015). Anatomical and functional evidence, such as spatially restricted cholinergic release (Disney & Higley, 2020; Parikh & Sarter, 2006), suggests that ACh can modulate specific cortical subnetworks. Yang et al. (Yang, Booth, & Zochowski, 2023) demonstrated that such spatial heterogeneity can drive synaptic reorganization and feature binding. In particular, regions with higher ACh concentrations (reflected as lower *g*_*ks*_ values) show enhanced excitability and plasticity, leading to localized network strengthening. Lohani et al. (Lohani et al., 2022) further showed that spatially distinct ACh signaling can dynamically regulate cortical states in behaviorally relevant ways. In future work, we aim to incorporate spatial heterogeneity of ACh into our model to better understand its role in DD.

### Comparison with other ACh models

Traditional ACh models used static modulation of HH-type conductance. More recent work, such as Codianni & Rubin (Codianni & Rubin, 2023), introduced dynamic models that simulate extracellular ACh concentration using differential equations, incorporating spike-triggered release, degradation, and receptor kinetics. Their model affects both M-current and synaptic input in real time, enabling feedback-based modulation of network states. Sajedin et al. (Sajedin, Menhaj, Vahabie, Panzeri, & Esteky, 2019) implemented a leaky integrate-and-fire (LIF) network to simulate cholinergic effects on orientation selectivity in visual cortex. Their model used *g*_*ks*_ modulation to emulate ACh’s role in sharpening tuning curves and reducing response variability. Our model, while simpler, is consistent with this framework: *g*_*ks*_ regulates ACh-like effects on excitability, synchronization, and deviance detection.

## Methods

### Neuronal model

The model includes a fast, inward Na^+^ current, a delayed rectifier K^+^ current, a leakage current, and a slow, low-threshold M-type K+ current. With membrane capacitance *C* = 1

*μF*/*cm*^2^, voltage *V*_*i*_ in millivolts, and time *t* in milliseconds, the current balance equation for the *i*-th cell is:

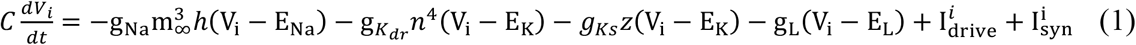

*I*_*drive*_ was a constant externally applied current. For the Na+ channel, activation was instantaneous, with the steady-state function:

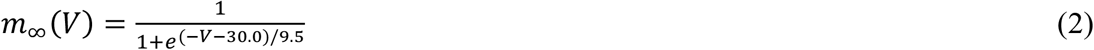

The inactivation gating variable *h*_*i*_ was described by:

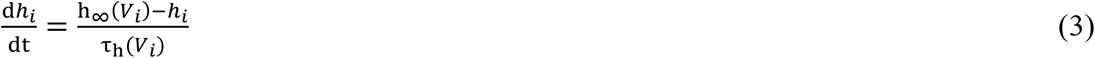

where 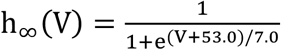 and 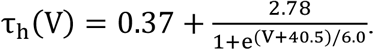

The delayed rectifier K^+^ current was gated by *n*_*i*_:

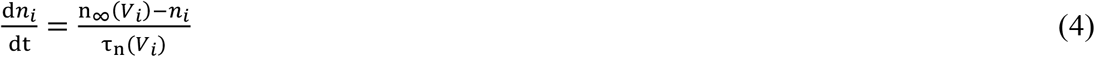

where 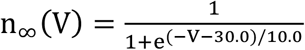 and 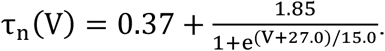

The slow, low threshold K^+^ current was gated by *z*_*i*_:

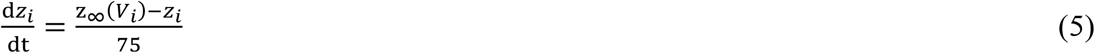

where 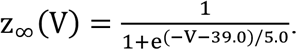

Other parameters were: *g*_*Na*_ = 24.0 *mS*/*cm*^2^, *g*_*kdr*_ = 3.0 *mS*/*cm*^2^, *g*_*L*_ = 0.02 *mS*/*cm*^2^, *E*_*Na*_ = 55.0 mV, *E*_*k*_ = −90.0 mV, and *E*_*L*_ = −60.0 mV. The maximal conductance of the slow K+ current 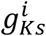 varied between 1.5 mS/cm^2^ (no ACh) and 0 mS/cm^2^ (strong ACh).

Synaptic interactions were modeled as transient changes in conductance.

### Synaptic dynamics

Synaptic currents were modeled using a conductance-based approach:

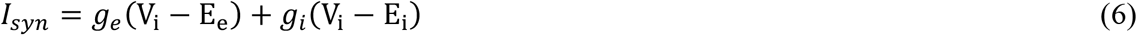

Here, E_e_ = 0 mV and E_i_ = −100 mV represent the excitatory and inhibitory reversal potentials, respectively. The synaptic conductances evolved as:

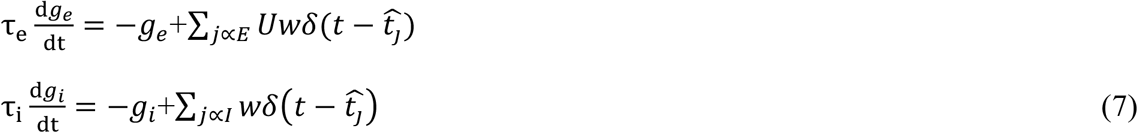

The time constants τ_e_ = 2 ms and τ_i_ = 4 ms emulate AMPA and GABA-A receptor dynamics, respectively. The synaptic weight w was fixed at 1 for all connections. The Dirac delta function δ(.) denotes a spike at time *t*_*j*_. The synaptic utilization factor U = 0.4 modeled the constant release probability at excitatory synapses.

### Short-term depression

STD was implemented for excitatory synapses only:

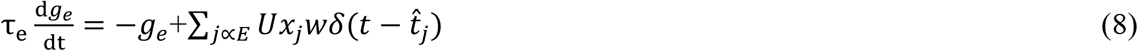

Here, *x*_j_ is a presynaptic depression variable defined per neuron:

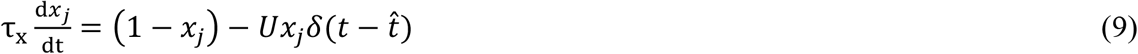

The recovery time constant was τ_x_ = 150 ms, and the release fraction was U = 0.4, consistent with auditory cortex data. This model excluded stochastic release, assuming deterministic transmission behavior.

### Network model

We simulated a cortical microcircuit consisting of 200 HH neurons placed randomly within a 2D circular region (radius = 4.0 mm). Neurons were 80% excitatory and 20% inhibitory.

Connectivity followed a distance-dependent probabilistic rule: each neuron projected to 10 random targets within a given radius (*r*_*exc*_ = 2 mm, *r*_*inh*_ = 1 mm for excitatory and inhibitory neurons, respectively). Conduction delays were neglected. Synaptic weights were fixed at 1 nS. Five network variants were generated using different random seeds.

### Stimulation setup

To study responses to stimuli with varying regularity, we used an oddball paradigm with five stimuli (A–E), each delivered as a 1.0 ms current pulse (12 µA/cm^2^). Each label mapped to a unique spatially localized neuron pair within an annular region (2.0–3.0 mm from center). Only the ensemble neurons received stimulation. Each trial began with one such pulse. This stimulus amplitude ensured at least one spike for any *g*_*ks*_ value, providing a consistent activation baseline.

### Oddball paradigm

We ran three types of oddball sequences (200 trials each, ISI = 100 ms): (1) Standard: A appeared frequently (p = 0.8), B was rare (p = 0.2); (2) Deviant: B was frequent (p = 0.8), A was rare (p = 0.2); and (3) Control: All five stimuli appeared equally (p = 0.2), serving as a neutral baseline. Stimulus blocks (5-element sequences) were randomly concatenated to form each condition. Each network ran four A/B pairings (1/2, 2/1, 3/5, 5/3) and one control sequence, giving 20 datasets total.

### Simulation details and code

Simulations used custom MATLAB scripts with forward Euler integration (0.01 ms timestep). At each step, synaptic currents, state variable derivatives, and synaptic conductance updates were computed. Each experimental block (standard, deviant, control) began from identical resting states, ensuring comparable dynamics. Trials were 100 ms long. We logged spike times for all neurons and constructed PSTHs (1 ms bins, 40 ms window post-stimulus A) to compare population responses across contexts.

### Deviance detection

Our primary quantity of interest was the stimulus-evoked response magnitude, denoted as R. We defined R as the average number of spikes fired in response to a specific stimulus across all relevant trials. This measure captures the mean trial-wise response and can be calculated at different levels of granularity.

We used subscripts and superscripts to clarify the response context. For example, 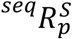 denotes the response of neuron(s) p, averaged across all trials of type S within the stimulus sequence seq. The index p could refer to: a specific neuron (e.g., *i, j*), a subpopulation (e.g., exc), or the entire network (if omitted), in which case R represents the median response across neurons. The trial type S could take one of the following values: A or B: indicating trials where stimulus A or B was presented, N: indicating non-target trials (stimuli C, D, or E). We defined the contrast in response between two conditions using the notation

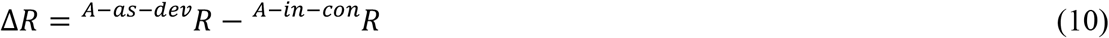

When aggregating over a population p, we used 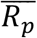 to denote the mean of individual responses within that population.

To isolate genuine DD, we followed established protocols that avoid conflating deviance with mere stimulus identity or adaptation effects. Specifically, we did not compare stimulus A in the A-as-dev and A-as-std sequences, as this would conflate deviance with adaptation due to frequent presentation. Instead, we compared responses to stimulus A in the A-as-dev and A-in-con sequences. The A-in-con condition serves as a neutral control, where all stimuli are presented with equal probability (p = 0.2). Thus, stimulus A appears as infrequently as in the deviant condition but without the context of an expected standard stimulus B. This control allows us to attribute any increase in response solely to statistical deviance, rather than repetition effects or physical differences between stimuli. To quantify this effect, we defined the Deviance Detection Index (DDI) as:

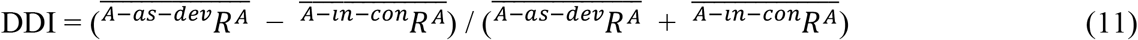

Here, 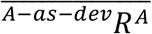 and 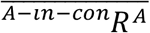 represent the mean number of spikes fired per neuron in response to stimulus A under the deviant and control conditions, respectively. A higher DDI value indicates stronger sensitivity to statistical deviance, independent of adaptation or identity effects.

### Spike frequency adaptation

Spikes were defined as upward membrane crossings of 0 mV. Only spikes after 100 ms (post-stimulus) were analyzed. ISIs were calculated from spike time differences. SFA was quantified as:

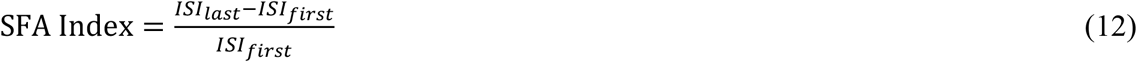

This index was computed only if ≥3 spikes occurred and *ISI*_*last*_ ≥ I*ISI*_*first*_. If spike rate increased (*ISI*_*last*_< *ISI*_*first*_), SFA was set to 0. Trials with <3 spikes were excluded.

### Mean phase coherence (MPC)

To compute phase 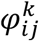 of spike *k* from neuron *i* relative to spikes of neuron *j*:

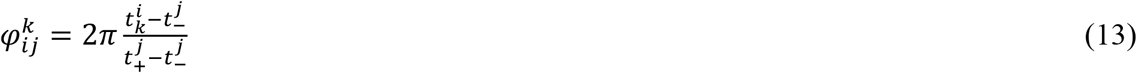

where 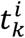 and 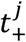 are the preceding and following spikes from neuron 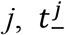 is the time of the last spike fired by neuron *j* before 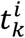. Pairwise coherence *r*_*i*j_ was computed as the magnitude of the mean complex vector:

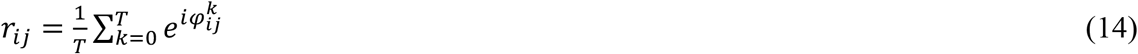

where T is the number of spikes fired by neuron *i* that are between a pair of spikes fired by neuron *j. r*_*ij*_ ranges from 0 (no synchrony) to 1 (perfect locking). Network-wide MPC was defined as the mean *r*_*ij*_ across all pairs where both neurons fired >30 spikes during the simulation.

## Data availability

All data used to produce the figures was generated via numerical simulations of the freely available code, which will be made publicly available upon acceptance.

## Code availability

The source code for our computational model and analysis scripts will be deposited in a public GitHub repository and archived on Zenodo with a permanent DOI upon acceptance of the manuscript. In the meantime, the code is available for review purposes upon request.

## Acknowledgements

We thank Felix B. Kern for helping with modeling. This work was supported by World Premier International Research Center Initiative (WPI), MEXT, Japan (to Z.C.C.); the National Key R&D Program of China (No. 2021ZD0201300 to Z.G.H.); the China Scholarship Council program (Project ID: 202406280171); and IRCN Retreat Brainstorming Awards FY2024 (No. 244700000001 to F. F.).

## Author contributions

Z.C.C. conceived and designed the research. Z.C.C. and F.F. designed the experimental protocol. F.F. built the models, performed numerical simulations, and analyzed the data. F.F. and Z.C.C. wrote the manuscript, F.F. prepared the figures, and Z.C.C. and Z.G.H. edited and revised the manuscript. All authors contributed to and approved the final paper.

## Supplementary Information

**Figure S1.**
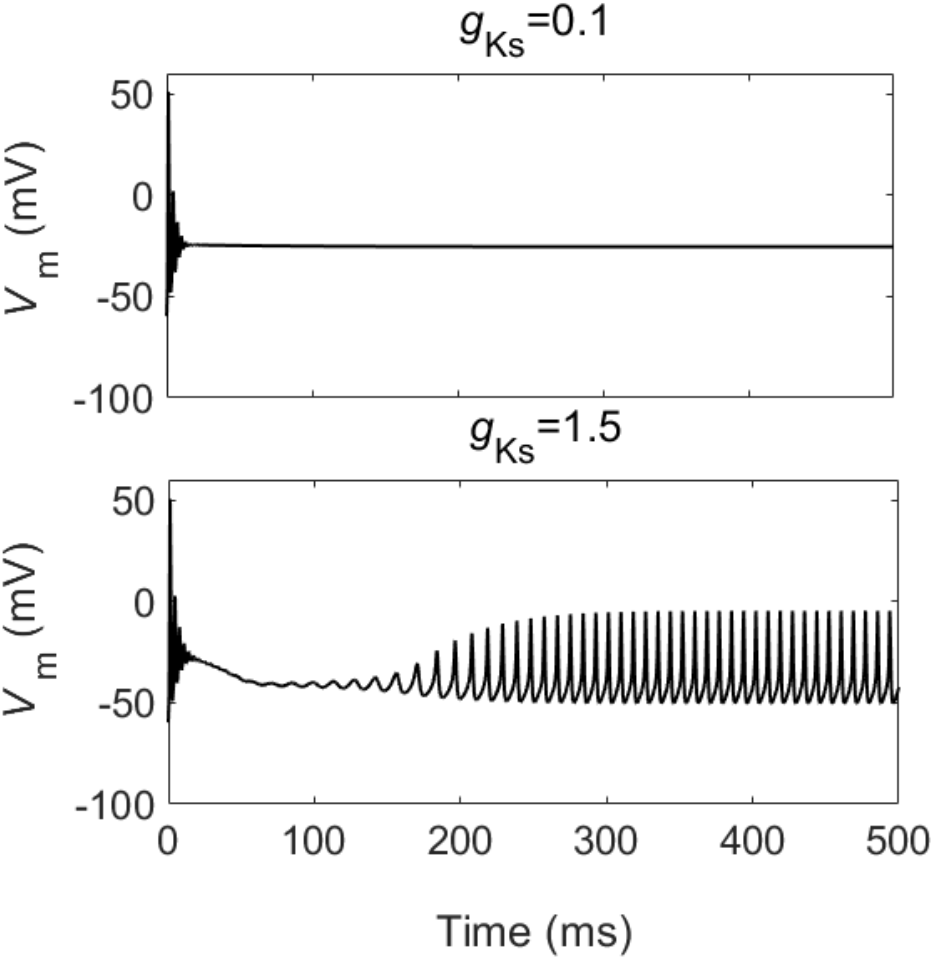
Evoked response under strong current. Change in membrane potential after an external stimulus current of 25 *μA*/*cm*^2^ under *g*_*ks*_ = 0.1 and 1.5.

**Figure S2.**
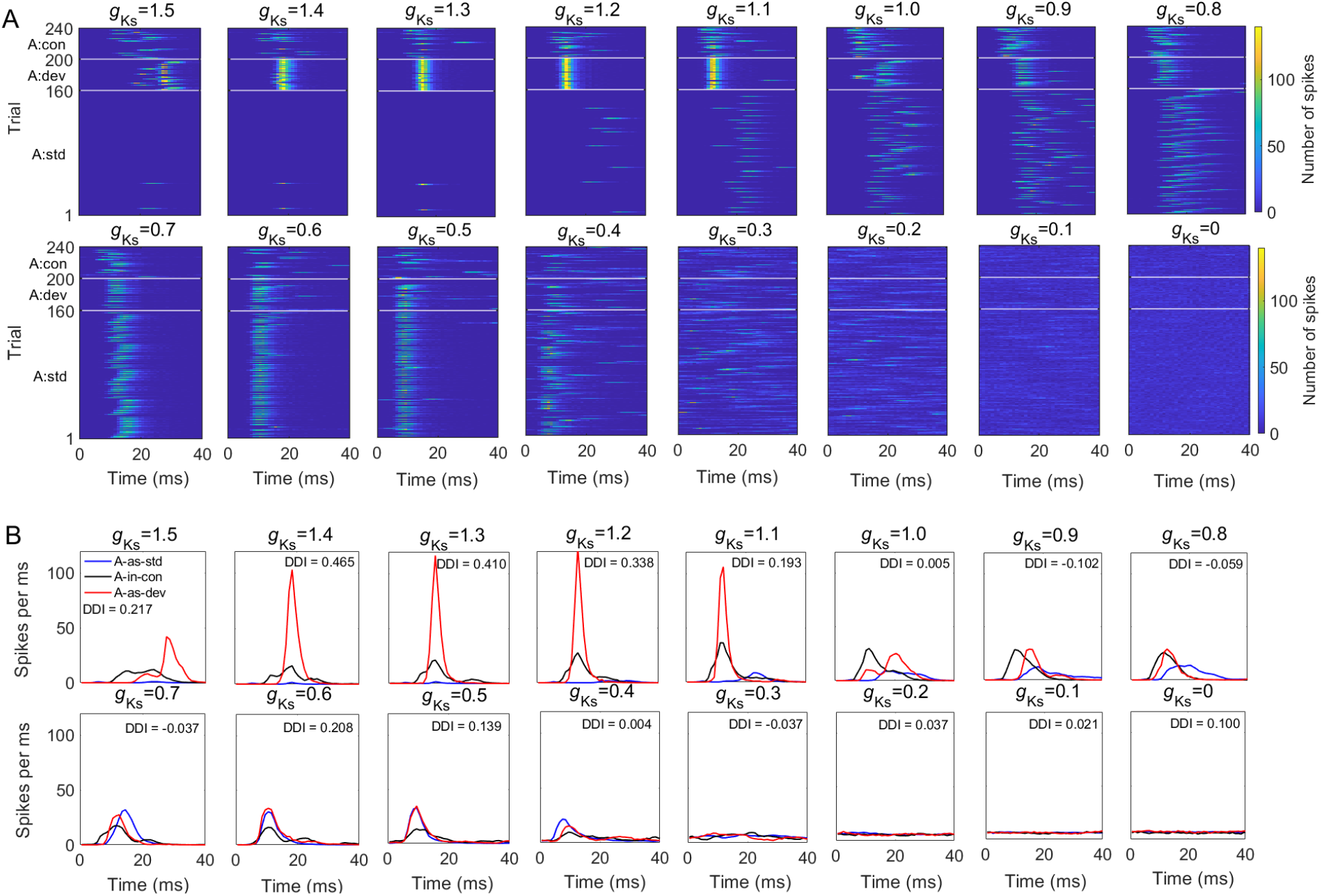
DD in a sample network with short-term depression. (**A**) Network responses to the target stimulus A. (**B**) Post-stimulus time histograms for network responses to stimulus A. Same representations are used as in Figure 4.

**Figure S3.**
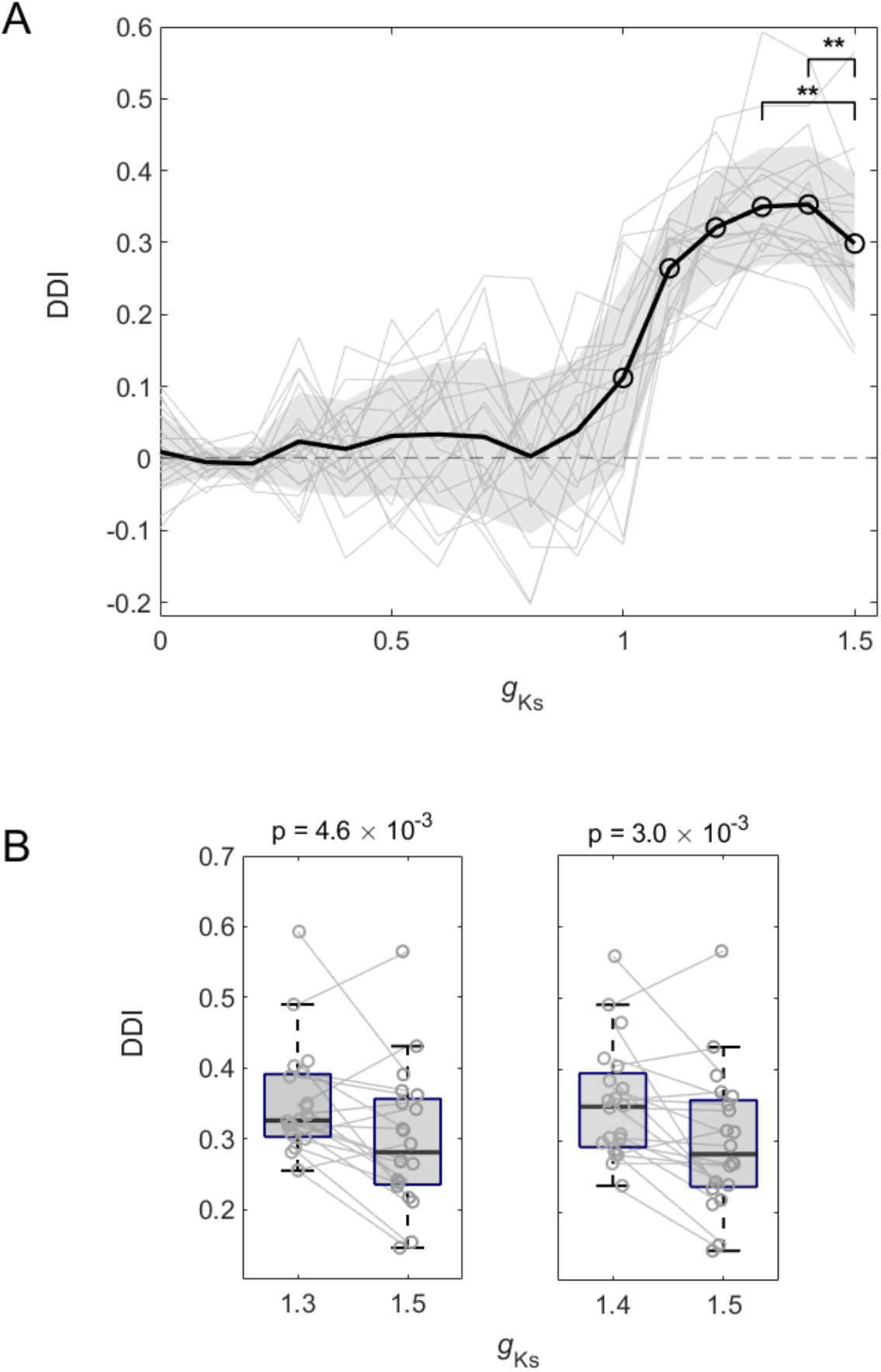
Cholinergic modulation of DD with short-term depression. (**A**) DDI values plotted against *g*_*ks*_ across 20 simulations (5 network configurations × 4 stimulation sites). (**B**) Conditions where DDI values (*g*_*ks*_ = 1.3, and 1.4) were significantly higher than at *g*_*ks*_ = 1.5. Same representations are used as in Figure 5.

**Figure S4.**
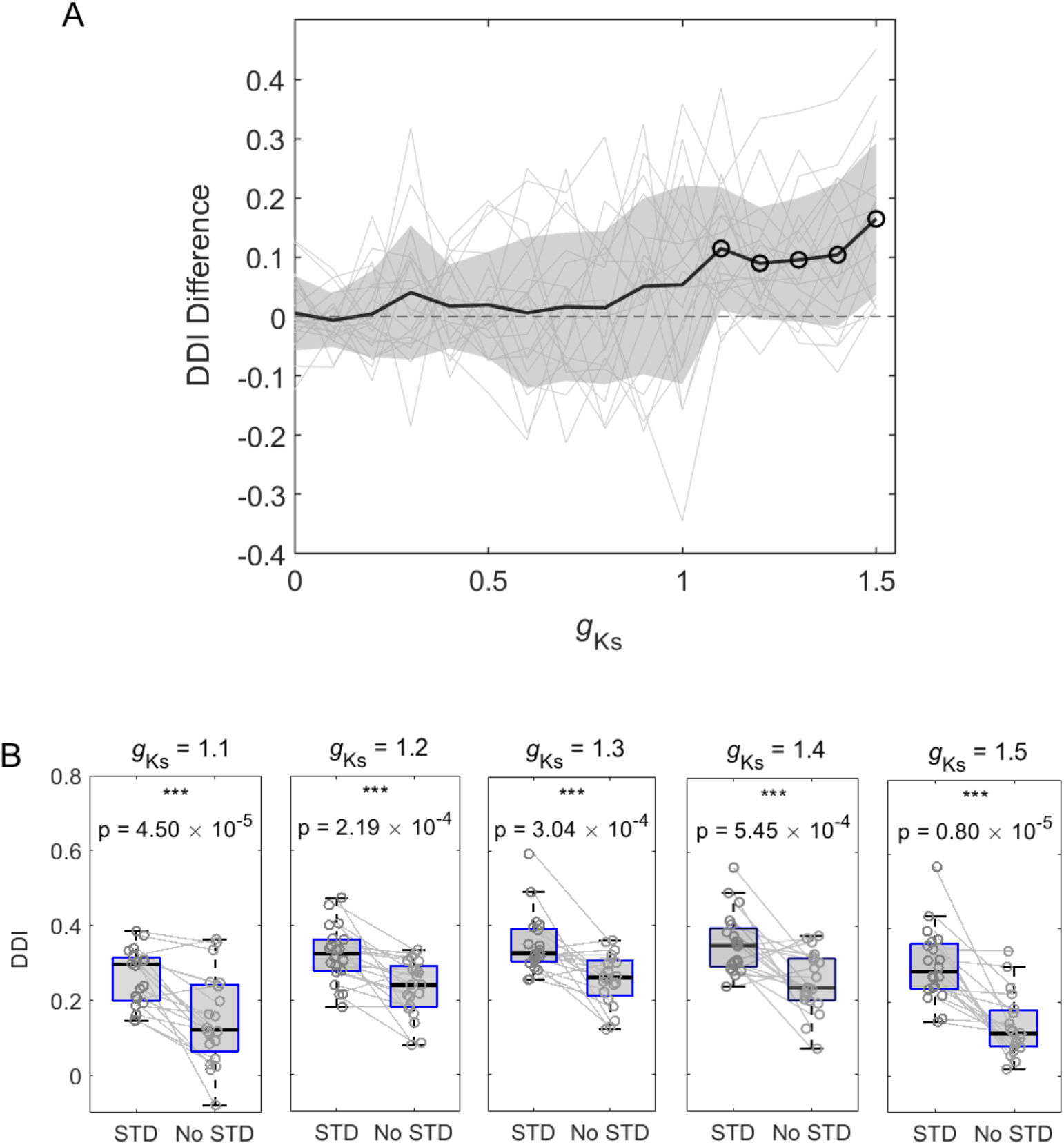
Comparison of DD with versus without short-term synaptic depression. (**A**) The difference in DDI values plotted against *g*_*ks*_ across 20 simulations (5 network configurations × 4 stimulation sites). (**B**) Conditions where DDI values (*g*_*ks*_ = 1.1, 1.2, 1.3, 1.4, and 1.5) were significantly higher than without short-term depression. Same representations are used as in Figure 5.

**Figure S5.**
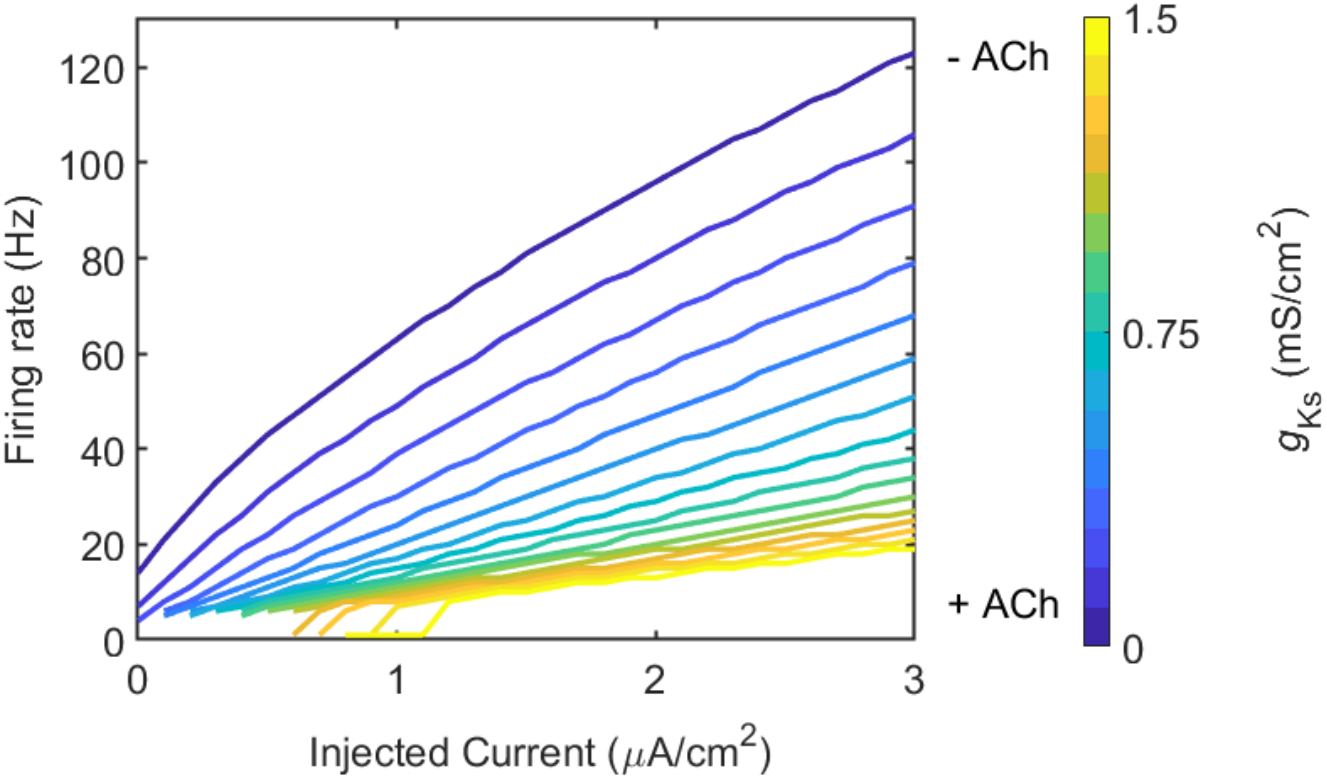
The f/I curves with different *g*_*Ks*_ values. Different colors represent the different *g*_*ks*_ values (shown as the color bar).

**Figure S6.**
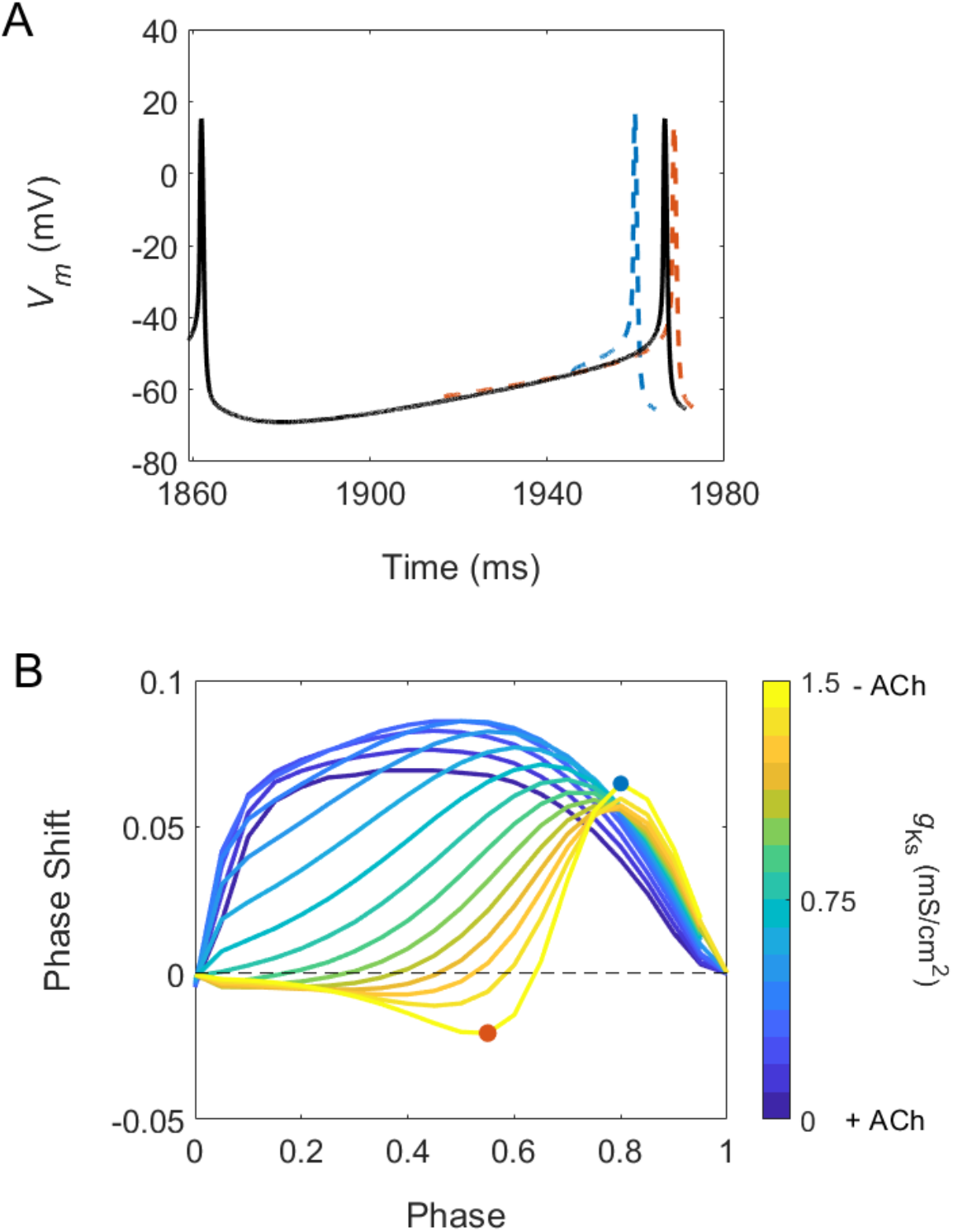
The phase-response curves with different *g*_*Ks*_ values. (**A**) The phase response curve is measured by comparing perturbed vs. unperturbed periods when neurons fire at a fixed frequency. When the next spike is earlier the phase response is positive (blue), when it is delayed it is negative (red). (**B**) Type 1 neurons have a strictly positive phase response curve (low ***g***_***Ks***_), while Type 2 neurons have a biphasic phase response curve (high ***g***_***Ks***_).

**Figure S7.**
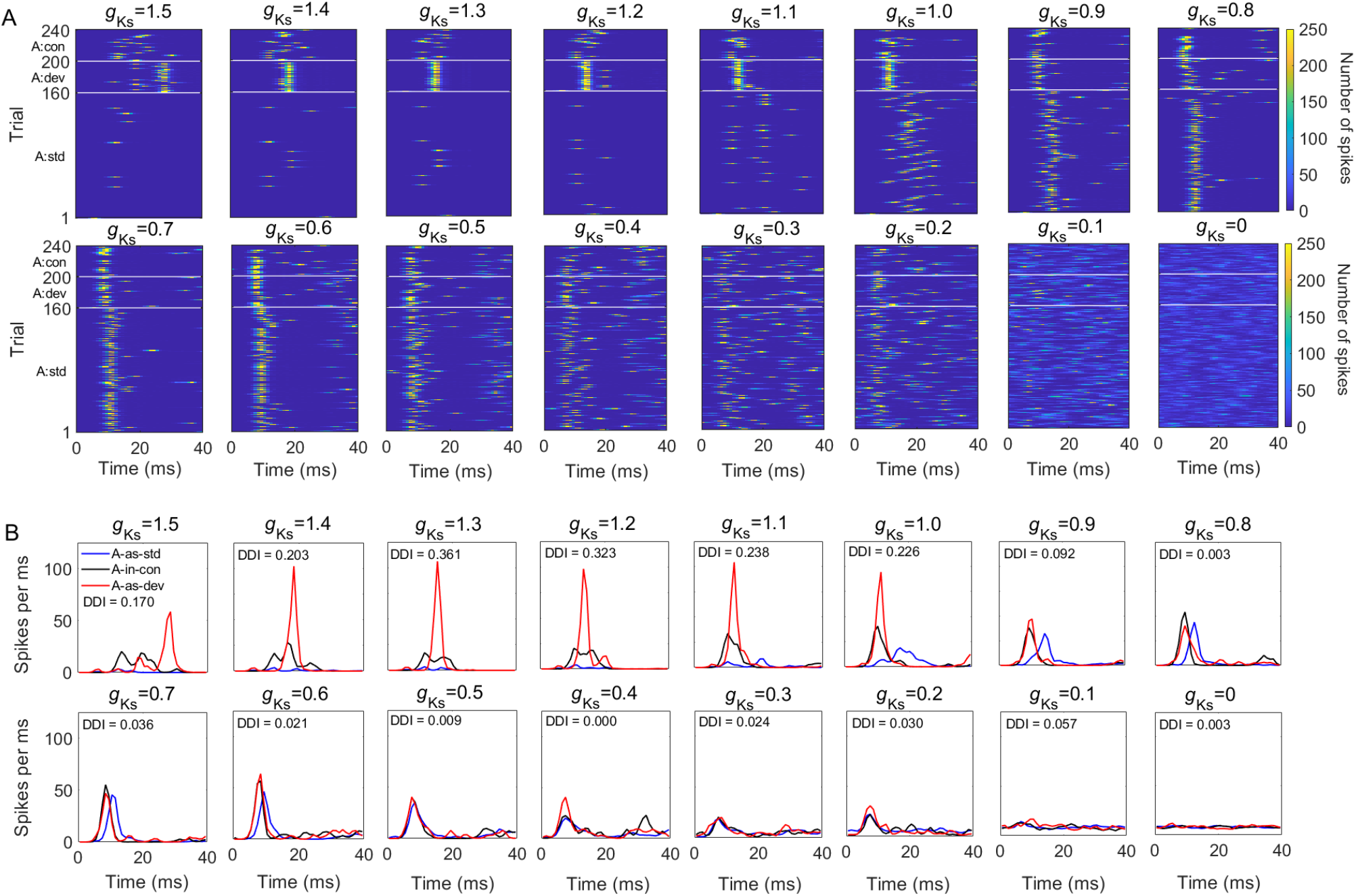
DD in the sample network under varying *g*_*ks*_ levels. (A) Network responses to the target stimulus A under three sequence types. A: std, A: dev, and A: con represent the evoked responses to stimulus A under the A-as-std (160 trials), A-as-dev (40 trials), and A-as-con (40 trials) sequences, respectively. Color indicates the total spike count across all neurons per millisecond. (B) Post-stimulus time histograms for network responses to stimulus A under the three sequence types. Sixteen *g*_*ks*_ conditions (1.5, 1.4, 1.3, 1.2, 1.1, 1.0, 0.9, 0.8, 0.7, 0.6, 0.5, 0.4, 0.3, 0.2, 0.1, and 0) are shown.

